# GW-CALL: Accurate Genome-Wide Variant Caller

**DOI:** 10.1101/079905

**Authors:** M. Ghareghani, S. A. Motahari, S. Khazaei, M. Tavassolipour

**Affiliations:** Department of Mathematical Sciences, Sharif University of Technology, P.O. Box 11365-8639, Tehran, Iran; Computer Engineering Department, Sharif University of Technology, P.O. Box 11365-8639, Tehran, Iran

## Abstract

The main challenge in reliable variant calling using DNA reads is to extract information from reads mappable to multiple locations on the reference genome. Conventional approaches ignore these reads and rely on reads mappable uniquely to the reference genome. These approaches fail to perform satisfactorily in variant calling within repeat regions which are abundant in many species including homo sapiens. This, in turn, lowers the reliability of any downstream analysis including poor performance in genome-wide association studies. GW-CALL, a fast and accurate variant caller, is proposed. GW-CALL exploits information of all reads in a genome-wide decision making process. In particular, it partitions the genome into several independent regions called clusters and incorporates an efficient algorithm to use all reads belonging to a cluster in calling variants within that cluster.

**Availability:** GW-CALL is implemented in C++ and is freely available at URL: brl.ce.sharif.edu/gwcall.

## 1 Introduction

With the advent of high-throughput next generation sequencing technologies, a large amount of sequencing data has been generated. The increasing amount of generated data has raised the demand for efficient computational tools to process these raw data. Generally, the problem of reconstructing a target genome using its reads, produced from a sequencing platform, is called DNA sequencing. The high level of similarity between DNA sequences of two individuals of one species enables us to make use of a sequenced genome of the same species as a reference genome in assembling the other genome. This problem is known as reference-based DNA sequencing, resequencing, or variant calling. It is called variant calling since with the availability of a reference genome, it suffices to call all variations between target and reference for a full reconstruction of the target sequence.

DNA sequencing has been applied to a wide range of biological problems, including disease association studies, discovering disease mechanisms, and phylogeny studies for finding the history of evolution. The accuracy of all downstream analyses depends on the way sequencing reads are processed reliably in the upstream. For a reliable association of genetic variations to phenotypes and disease, as an example, it is essential to be able to call variants accurately.

In cancer genomics, studying of somatic variants plays an important role in finding causal factors of a particular cancer [9], [5]. Many variant callers are designed to detect variations in cancer genomes including VarScan [6], [7] and MuTect [2].

Due to evolutionary reasons, the genome sequences of individuals contain repetitive paralogous regions [17]. Redundant genomic regions are the main source of ambiguity in both read mapping and variant calling [18]. Indeed, alignment errors in these regions significantly affects the downstream task of variant calling. A recent study focussing on challenges of calling repeat regions has revealed that common variant callers GATK [14] and samtools mpileup [10] have poor performance in repeat regions [3].

The standard approach for resequencing is mapping-based variant calling in which all reads are mapped to the reference genome and then the target genome is reconstructed using the information of mapped reads. The key challenge in repetitive regions is that their reads can be mapped to multiple locations on the genome, and one cannot distinguish their original positions, i.e., the positions where they have been sampled from. A number of strategies have been employed in order to evade the problem of alignment ambiguity, such as discarding all multi-reads (multiply-mapped reads) and utilizing only unique-reads (uniquely-mapped reads) in variant calling [13, 12], or selecting the best mapping position, for example the position with lowest hamming distance with each read. All such strategies can be shown to produce erroneous calling in some cases.

The majority of mapping-based callers, including GATK [14] and samtools mpileup [10], do not deal with the complexity of repeat regions and make the simplifying assumption that there is no alignment error, i.e., all reads are mapped to their correct position. Based on this presumption, they locally call each genomic locus by simple genotyping models using the information of the pileup data in that position. As a result, they fail to call repetitive parts of the genome accurately. Sniper [15] is a mapping-based caller that considers ambiguity of multi-reads in a more realistic genotyping model. In fact, it performs a more global variant calling in repeat regions where ambiguous reads are mapped to the genome. However, the running time of its calling algorithm grows exponentially with the length of repetitive regions, making it nearly impractical for genomes with many repetitive elements, such as the human genome.

Some genomic parts, namely repeat regions, are more complicated and ambiguous for calling, but the remaining loci are accessible and can be called confidently. We use a heuristic approach to differentiate between accessible and ambiguous genomic loci. Our results show that ambiguous positions cover only a small fraction of genome. This fact inspired us to design a variant caller, which we refer to as GW-CALL, that calls all accessible loci by a rather simple algorithm, and then process the remaining ambiguous parts, called critical points, in a more computationally heavy, but still practical, procedure.

The ambiguity of multi-reads creates dependency between critical points, i.e., variant calling in a critical point influences variant calling in other critical points, in the presence of multi-reads. We refer to the example in section 3.1 for a detailed discussion. This fact makes the task of variant calling in repeat regions complicated and makes us call variants in critical points collectively. We cluster critical points such that those critical points belonging to one repeat region are put into the same cluster. The critical points in a cluster are connected to each other, but they are independent from the critical points in other clusters. Therefore, we can perform variant calling for critical points in each cluster independently.

We use a heuristic approach in order to decrease the number of candidate target sequences for variant calling in a cluster. To this aim, we partition each cluster into several chunks. Every chunk contains those critical points of a cluster which are concentrated in one part of the genome. For each chunk, we select a list of most probable candidate sequences for reconstructing target genome. We utilize a dynamic programming algorithm for efficiently computing these lists for chunks. Finally, for each cluster, we perform a global variant calling procedure in which we find the most probable selection from the lists of candidate sequences of its chunks.

## 2 Methods

We use the following notation: If *S* is a string, then *|S|* represents the length of the string and *S*[*χ*] represents a subsequence of *S* indexed by the set of locations *χ*. In particular, *S*[*i*] for *i* ∈ {1, …, |*S*|} is the *i*th letter of *S*.

Let the reference genome *X* be a string of length *n* over the alphabet Γ = {A, C, G, T}, and a target genome 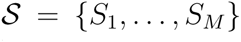 consists of *M* DNA sequences of length *n* (*M* represents the polyploidy number of the target genome). Variations between reference and target genomes are assumed to be Single Nucleotide Variations (SNVs). Suppose every target genome has SNV probability *ϵ*, independent of other sequences; that is, for all *i* ∈ {1, …, *M*} and *j* ∈ {1, …, *n*};

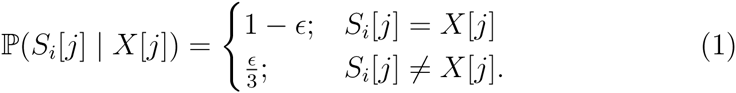

For brevity, in the case of haploid targets, we denote the target sequence by 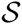. We have a set of reads 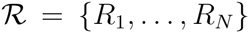, each one of length *L*, extracted from one of the target genomes. The extraction is assumed to be distributed uniformly across the genomes. We also assume that the process of extracting reads is noisy in which a base at the *q*th position of the read is read erroneously with probability *p*_*q*_, where 1 ≤ *q* ≤ *L*. More precisely, if *R*_*j*_ is sampled from location *d* of the genome *S*_*i*_, we have: for all *q* ∈ {1, …, *L*};

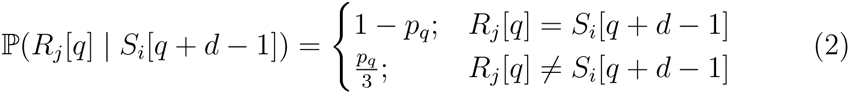

The variant calling problem is the process of target reconstruction, given the reference genome and the set of reads. It consists of two distinct problems: haplotype phasing and genotyping. Haplotype phasing amounts to obtaining the target genome 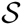 with all chromosomes phased. In genotyping, we aim to reconstruct the target genotype 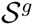, given *X* and 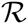. Here, 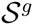 determines the number of occurrences of alphabet letters in all locations without specifying which base belongs to which chromosome. Hence, 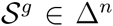, in which Δ is the following set:

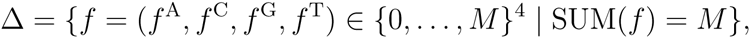

In other words, if the *i*th location of 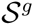 is 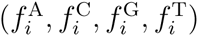, it means that there are 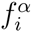 chromosomes with alphabet *α* in the *i*th location of the target.

Our approach is a mapping-based variant caller that takes as input a reference genome and a mapped file including information of mapping reads to the reference (current implementation accepts SAM file [11]), and returns an estimate of the target sequence. We assume that the mapper is nearly perfect, i.e., the true location of a read is almost always within the list of reported positions of that read. It is worth mentioning that a dummy mapper that reports for each read all the locations of the genome is a perfect mapper. However, a perfect mapper which has minimal list sizes is favourable as it hugely reduces the computational complexity of the variant calling.

### 2.1 Key observation and basic tools

We first present an example to show the limitations of local variant calling and to see the out-performance of global decision making. We then present a new MAP approximation which is used in our variant calling algorithm. We also review some basic tools that we utilize in variant calling.

#### 2.1.1 Global decision making

Let us recall two commonplace mapping-based scenarios: **unique-map** and **all-map**. We then discuss their limitations in calling variants in repeat regions. In the unique-map strategy, all multi-reads are discarded, and the variant calling is performed solely based on unique-reads. Discarding multi-reads leads to losing sequencing data in repetitive regions. Therefore, in this scenario, we miss all variants occurred in these parts.

The all-map strategy utilizes all reads for variant calling by a simple variant calling method. In other words, it locally calls each locus based on its pileup data, regardless of calling information of other loci. In this manner, incorrect mapping of reads in repeat regions may lead to erroneous calling in some positions.

Figure 1 shows that simple variant calling methods are not powerful enough for accurate calling in the presence of ambiguous multi-reads. The figure represents a haploid genome with three repetitive parts. We assume three noiseless reads are sampled from each of these parts. If we map all reads to the reference genome with hamming distance at most two, the reads sampled from each of these regions map to all of these parts. The unique-map strategy removes all of these reads and hence cannot call the SNV in location *y*_6_. The all-map strategy makes erroneous calling in positions *y*_4_ and *y*_5_. It calls these positions by C and A respectively, due to its local decision making. In fact, existence of erroneously mapped multi-reads covering these positions leads to defective calling.

**Figure 1:**
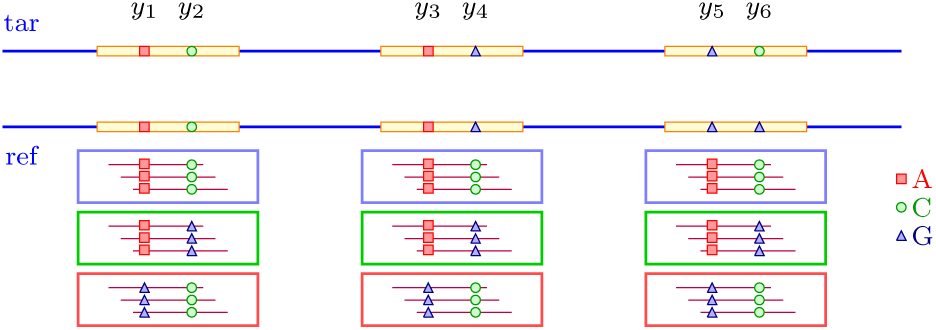
An example of read mapping in a repeat region with three repetitive elements. Reads are assumed to be noiseless. *y*_6_ is the variant between the target and reference genomes. The unique-map strategy discard all reads and looses the coverage. The all-map strategy makes false decision in *y*_4_ and *y*_5_. From information embedded in reads AC, AG, and GC are the only possible pairs at each locus. A global decision maker can find the best permutation of these pairs.

In a more global view, we realize that we should call the pairs of positions (*y*_1_*, y*_2_), (*y*_3_*, y*_4_), and (*y*_5_*, y*_6_) by values AC, AG, and GC, and we ought to use all of these values since reads are noiseless. Therefore we should select a permutation of these values for target reconstruction in these three genomic regions. The most likely permutation is the one with the lowest hamming distance with the reference genome, which yields the original target sequence. As a result, we can reconstruct the target sequence without any false detections, by using the information of multi-reads and making a global calling.

This example has motivated us to develop a step-by-step procedure for calling repeat regions with high accuracy. First, we call all accessible parts of the genome and mark the ambiguous loci (e.g., positions {*y*_1_, *y*_2_, *y*_3_, *y*_4_, *y*_5_, *y*_6_} in Figure 1.) for further processing. Second, we partition the set of ambiguous points into units called chunks, based on their connectivity created by multi-reads. We then obtain a set of candidate values for target sequence corresponding to each chunk. Finally, we make a global selection of candidate sequences for all chunks belonging to a repeat region.

#### 2.1.2 A new MAP approximation

We present a new MAP approximation which will be used as a basis in our variant calling algorithm. We propose some preliminary definitions and then present a basic proposition for MAP estimation.

##### Definition 2.1 (mapping interval and mapping area of a read)

*Let R be a read mapped to locations* {*x*_1_, …, *x*_*d*_}. *Every set* {*x*_i_, *x*_*i*_ + 1, …, *x*_*i*_ + *L* − 1}, *for* 1 ≤ *i* ≤ *d, is called a mapping interval of R. The union of all mapping intervals of R is called the mapping area of R, denoted by MP* (*R*).

Let χ ⊆ {1, …, *n*} be a subset of genomic locations. We denote by 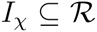 the set of all reads that each of which covers at least one location in *χ*. More precisely, *I*_*χ*_ is defined as follows:

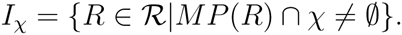

Let *X*[*χ*] and 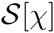 be reference and the target sequences corresponding to *χ*, respectively. Suppose 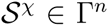 is a sequence that is equal to 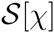 in *χ* and equal to the reference genome in other genomic locations. The MAP rule reveals an estimate 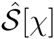 of the target sequence such that:

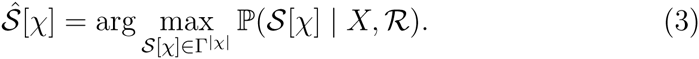

##### Proposition 2.2

*The posteriori probability* 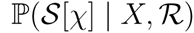 *is approximately proportional to the following term*:

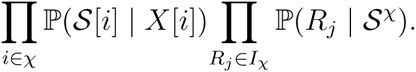

In the Supplementary Material, the reader can find a proof of the proposition.

Let us denote the logarithm of the expression in the proposition 2.2 by 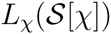. We define function 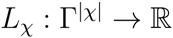 as bellow:

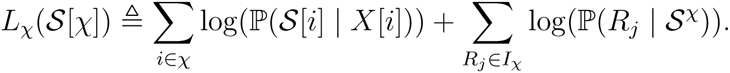

Therefore, for a subset *χ* of our interest, our goal is to find a genotype 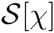 that maximizes 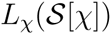. For a singleton set *χ* = {*i*}, we use the simplified notation *L*_*i*_ instead of *L*_{*i*}_.

#### 2.1.3 Other tools

We use two basic tools in our variant caller: Viterbi algorithm [19] and the junction tree algorithm [1]. In fact, we use a Viterbi-like dynamic programming algorithm for finding a set of candidate sequences for subsets of genomic locations called chunks. We also use the junction tree algorithm for global variant calling in each repeat region of the genome. We detail our approach, including an illustration of how we use these tools, in the subsequent section.

### 2.2 Detailed description of GW-CALL

Since the diploid case is a simple extension of the haploid case, we present our algorithm for the simpler case of haploid genome. The general case is explained in section 2.3.

Our variant caller, GW-CALL, consists of six steps. In the first step, we utilize a heuristic method to distinguish between accessible and ambiguous parts of the genome. We call all accessible loci in this step and mark the rest as critical points. The critical points —which mostly lie on the redundant genomic regions— are the most difficult parts for calling and, at the same time, cover a very small portion of the genome. Hence, it makes sense to call them in a more expensive procedure, in the next stages, in order to improve their calling accuracy.

In the second step, we partition the set of critical points into independent sets called clusters. In the third step, we partition each cluster into different sets called chunks, such that every chunk is a clump of critical points in a cluster. Then, we create a graph in which each connected component corresponds to a cluster in the forth step. For performing a global variant calling in each cluster, we find a list of most probable sequences for every chunk by an efficient dynamic programming algorithm, in the fifth step. Finally, in the sixth step, we search for the most probable selection from the lists of candidate sequences of chunks in every cluster. We present a detailed explanation of the steps of our algorithm in the rest of this section.

#### Step 1. Calling accessible loci and finding critical points

Let *i* ∈ {1, …, *n*} and *u*_*i*_, *v*_*i*_ ∈ Γ be the two alphabet letters with the first and second maximum values of 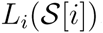. We define a calling confidence measure *C*(*i*) for location *i* as follows:

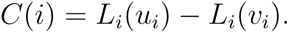

Intuitively, low calling confidence shows ambiguity in variant calling. Consider constant real numbers *τ* ≥ 0 and 0 < *ρ* ≤ 1. We call location *i* by base *u*_*i*_, if at least one of the conditions **A**, **B**, or **C** holds. Otherwise, we mark this position as a critical point.

A. *I*_{*i*}_ is empty (i.e., there is no coverage in position *i*),
B. All reads in *I*_{*i*}_ are unique,
C. It holds that *u*_*i*_ = *X*[*i*] (i.e., the base *X*[*i*] itself has the maximum posteriori probability) and at least one of the following conditions holds:
  - *C*(*i*) > *τ*,
  - The ratio of multi-reads mapped to this locus is less than *ρ*.

In the rest of this paper, *χ* denotes the set of all critical points.

#### Step 2. Partitioning critical points into clusters

We cluster critical points based on their dependencies created by multi-reads. We present an exact definition of clusters and explain how the problem of calling critical points in a genome can be divided into subproblems of calling critical points in each cluster separately.

We say two critical points are *connected*, if they are both located in the mapping area of a read. We say that a set *A* ⊆ *χ* has *connection property*, if for every *y, y′* ∈ *A*, there is a sequence (*y* = *y*_0_, *y*_1_, *…, y*_*d*_ = *y′*) of critical points such that for every 1 ≤ *i* ≤ *d*, *y*_*i*−1_ and *y*_*i*_ are connected.

Partition *χ* into **clusters** *χ*_1_, …, *χ*_*Q*_ such that each *χ*_*i*_, for 1 ≤ *i* ≤ *Q*, is a maximal subset of *χ* with connection property. Obviously, this partitioning is unique. It can be easily shown that the objective function *L*_*χ*_ is factorized as follows:

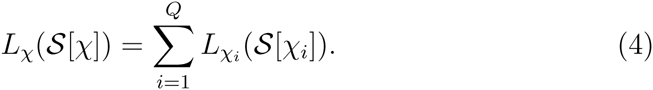

As the domains of functions *L*_*χi*_, for 1 ≤ *i* ≤ *Q*, are mutually disjoint, we can maximize these functions separately for maximizing *L*_*χ*_.

#### Step 3. Partitioning clusters into chunks

The critical points of a cluster can be dispersed in different parts of the genome. Thus, we partition every cluster into different blocks, called chunks. Each chunk is a clump of critical points of cluster in one part of the genome, which are connected by multi-reads. More precisely, we define chunks as follows:

##### Definition 2.3 (chunk)

*Assume that critical points of the cluster χ*_*i*_ ⊆ *χ are sorted. Each chunk of cluster χ*_*i*_ *is defined to be a maximal set of consecutive critical points in χ*_*i*_, *in which every two successive critical points are located in a mapping interval of a read.*

#### Step 4. Constructing adjacency graph

Creating chunks and clusters can be performed using a simple algorithm. We first scan the set of sorted critical points to find which successive critical points are put into the same chunk, so we can construct the set of all chunks in the genome. Then, we create an undirected graph *G* = (*V, E*), in which *V* denotes the set of all chunks, and {*C, C′*} ∈ *E* if and only if *C* and *C′* are adjacent. Here, we call two chunks *C* and *C′* adjacent, if the sets *I*_*C*_ and *I*_*C′*_ have a read in common. Every connected component in this graph corresponds to a cluster. We refer to the corresponding component to each cluster as an adjacency graph. An example of critical points, chunks and clusters in a genome is presented in the Figure 1 in Supplementary Material.

#### Step 5. List decoding for chunks

A list decoder is proposed to obtain a list of candidate sequences for each chunk. Conceptually, it amounts to collapsing all the critical points of a chunk into a few possible combinations. To make it more clear, let us consider a given chunk *C* consisting of *p* critical points. These critical points can potentially take all 4^*p*^ possible combinations of the four letters {*A, C, G, T*}. The list decoder finds only *k* (*k* can be as low as 2) possible combinations out of 4^*p*^ possible ones. The detail of the list decoder and all the derivations are presented in Algorithm 1 in Supplementary Materials. Here, we only presents the main ideas behind the proposed list decoder.

The objective function used to obtain the *k* best combinations is similar to the one presented in Section 2.1.2. Naively, we need to compute an approximate to MAP at 4^*p*^ different points and obtain the *k* highest ones. This is computationally intensive and not feasible for even moderate values of *p*. However, the objective function in our problem can be partitioned and maximized efficiently using a dynamic programming algorithm similar to the Viterbi Algorithm [19].

The list decoder first splits the chunk *C* = {*y*_1_, *y*_2_, …, *y*_*p*_} into overlapping segments such that each segment contains minimum number of critical points and no read belongs to more than one segment. There is an efficient algorithm to find these segments. Moreover, it can be shown that this split is unique. An illustrative example is given in Figure 2. In this way, the objective function can be written as the sum of functions over critical points of segments. However, it does not imply that each function can be optimized separately as there exit critical points common between these functions. Interestingly, we can employ a dynamic programming algorithm that efficiently resolves the problem.

**Figure 2:**
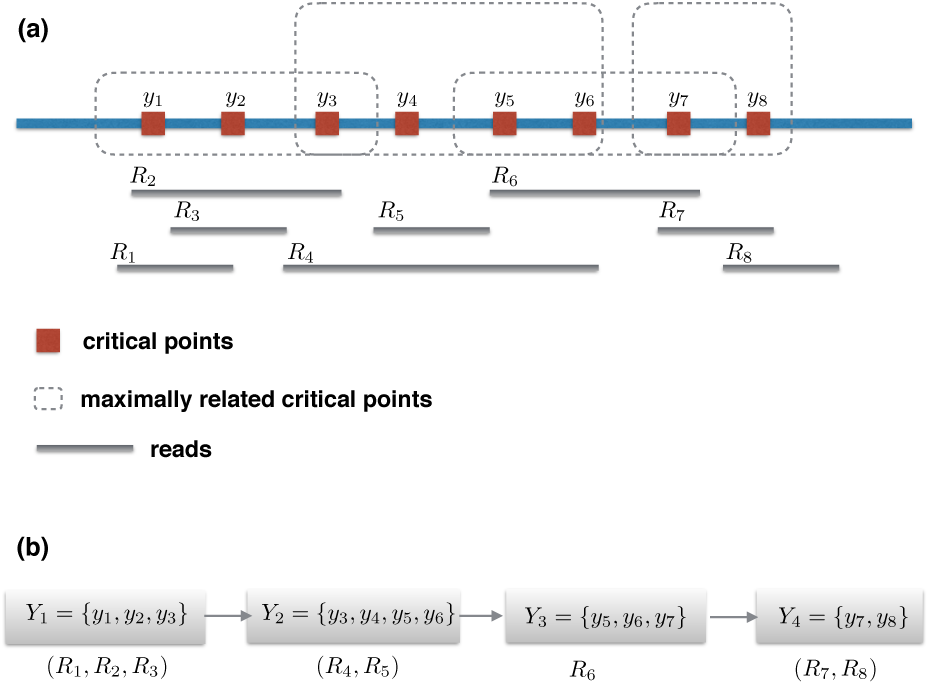
List decoding. (a) The critical points {*y*_1_, …, *y*_8_} are segmented into maximally related critical points incited by dashed boxes. (b) Critical points within a read is included in one of the segments. Reads are assigned to the segments and state space diagram is created, indicated at the bottom of the figure.

Even with the separation of the objective function, the critical points within a segment need to be processed jointly, i.e., the sate space of the dynamic program depends exponentially on the number of critical points within a segment. To overcome this, we limit the number of possible combinations of these critical points to the cases where the hamming distance between reference and target within the segment is bounded by some constant *h*. In this way, the possible number of states does relate polynomially to the number of critical points.

The dynamic programming algorithm consists of a forward and a backward procedure. In the forward procedure, we first compute the objective function for the first segments. Then, for every segment, we find the most *k* probable paths from the previous segment and store these paths in a collection of pointers called *P*. Finally, for each of *k* best paths between the first and last segments, we store a pair including the corresponding final decoded segment and the index of the path in the *k*-tuple list of that state in a set called *E*. The sets *P* and *E* are returned as the output of the forward procedure. The backward procedure uses these outputs and backtracks from the *k* best pairs in *E* following the stored paths in *P* to reconstruct the *k* most probable sequences.

After each run of the dynamic programming algorithm, we obtain a list of candidate sequences for each chunk. If we select the first element of the list for each chunk, we obtain a new target sequence. We can replace the reference genome with this new target sequence and run the dynamic programming algorithm again to obtain new lists of candidate sequences for chunks and repeat this procedure. Therefore, we can run the dynamic programming algorithm in an arbitrary number of iterations. We experimentally observed that two iterations significantly improves the performance of our variant caller.

#### Step 6. Calling chunks in a cluster

Consider a cluster *χ*_*i*_ ⊆ *χ* in which each chunk has *k* candidate sequences, computed in the aforementioned list decoding algorithm. We denote the number of chunks in *χ*_*i*_ by *n*(*χ*_*i*_). We can view every chunk as a set of variables taking values in the set {1, …, *k*} and view *L*_*χ*_*i*__(*·*) as a function of these variables. In other words, every configuration of {1, …, *k*}^*n*(*χ*_*i*_)^ relate to a selection of sequences from the list of chunks in *χ*_*i*_ and hence determines the value of target bases corresponding to *χ*_*i*_; so, it determines the value of 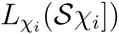. Our final goal is to find a configuration from the set {1, …, *k*}^*n*(*χ*_*i*_)^ that maximizes the function *L*_*χ*_*i*__(*·*).

A naive solution to this problem is to perform an exhaustive search in the whole possible configurations of chunks in {1, …, *k*}^*n*(*χ*_*i*_)^, which is not a practical solution for large clusters. As an alternative, we can employ a sort of dynamic programming algorithm called the junction tree algorithm [1] for maximizing *L*_*χ*_*i*__(*·*). The running time of this algorithm is exponential in the treewidth of the adjacency graph of the cluster. Therefore, we can utilize it for large cluster with small treewidth to make it tractable. Our simulation results show that there are a lot of large clusters with small treewidth. Note that, we have used algorithms presented in [4, 16] for construction of the junction tree.

In summary, if the treewidth of the adjacency graph of a cluster is reasonably small, we run the junction tree algorithm. Otherwise, we select the first candidate sequence of each of its chunks for reconstructing the target genome corresponding to this cluster.

#### 2.3 Variant calling in diploid genomes

We discuss how our variant calling algorithm can be modified to handle the diploid genomes. In the first step of the algorithm, we should use the alphabet set Δ instead of Γ. There is a tiny difference in computing the probability of reads in the diploid case which is explained in Supplementary Materials.

For the list decoding task, we use the alphabet set Γ^2^ and define a new state space, corresponding to the new alphabet, for critical points of a chunk. With this new alphabet set, every path through the states of critical points of a chunk correspond to two target haplotypes. Note that for some of these paths, there is a complementary path that corresponds to the same target haplotypes with different order. Since the order of haplotypes is not important, we should choose one path among each pair of complementary paths in the list decoding task.

If we replace the target haplotypes (*S*_1_, *S*_2_) with the sequence *S* in the function *L*_*C*_(*·*), the objective function factorizes similar to the haploid case. Therefore, we can use a similar dynamic programming algorithm to find the set of *k* most probable paths through the state space of a chunk. In the final step of the algorithm, we search through different selections from the list of chunks in a cluster and find the most probable one. The algorithm for this step resembles the haploid case.

## 3 Results

GW-CALL is implemented in C++ and its performance is evaluated on several data-sets obtained from human chromosome 19. Human Build 38 patch release 5 (GRCh38.p5) is used as the reference genome. The reference genome is also trimmed by removing all ambiguous characters (N bases).

The target genomes are obtained from the reference genome as follows. Given the SNV probability (*ϵ*), each base on the reference genome is either remained unchanged with probability 1 − *ϵ* or randomly mutated to another base with probability *ϵ*. In this way, a target genome is obtained. To generate multiple genome, this process is executed independently for each target genome.

Reads are obtained from the target genomes as follows. Given the read Length (*L*), the read error probabilities (*p*_*q*_’s) which is assumed to be the same and equal to *p* over all bases, and the coverage depth (*c*), reads starting points are obtained from a point process with independent inter-arrival time which are assumed to follow a geometric distribution with parameter *cL*. Each base in a read is either exactly the corresponding base on the target genome with probability 1 − *p*, or randomly changed to a new base with probability *p*.

After generating reads and target genomes, reads are mapped to the reference genome by mismatch 3 in all simulations using Bowtie [8]. Reads that are mapped to more than 30 locations on the reference genome are removed and the rest of reads are used for variant calling. In the current implementation of GW-CALL, we only call SNVs and short indels are not considered. Therefore, all reads which are mapped to the reference genome with indels are also discarded.

GW-CALL requires several adjusting parameters to be specified by the user. The default parameters that are used in the experiments are as follows: *τ* = *κ* = 5, *ρ* = 0.5, *h* = 2 and the list decoder size *k* = 2. GW-CALL is iterated twice in all the experiments. To reduce the complexity of the junction tree algorithm, we remove all junction trees with maximum clique size greater than 20.

GW-CALL is compared with the unique-map local strategy which is implemented in many variant calling tools. In this strategy, reads that can be mapped uniquely to the reference genome are only considered for variant calling and each base is called based on all the bases covering it. As a reference point to our results, we have also considered a perfect caller. In this caller, aided by a genie, all reads are anchored correctly to their location and best decision making is made for calling variants. Note that this variant caller can be counted as a lower bound on the error probability of any real variant callers as extra information is provided to the caller.

In Figure 3, we have compared the three variant callers for a set of different read lengths, *L*, ranging from 50 to 100 with step size 10. Other parameters are fixed with values: *p* = 0.01, *c* = 20, and *ϵ* = 0.001. The result shows that GW-CALL performs very close to the optimum caller while the unique-map caller performs poorly when the read length is short.

**Figure 3:**
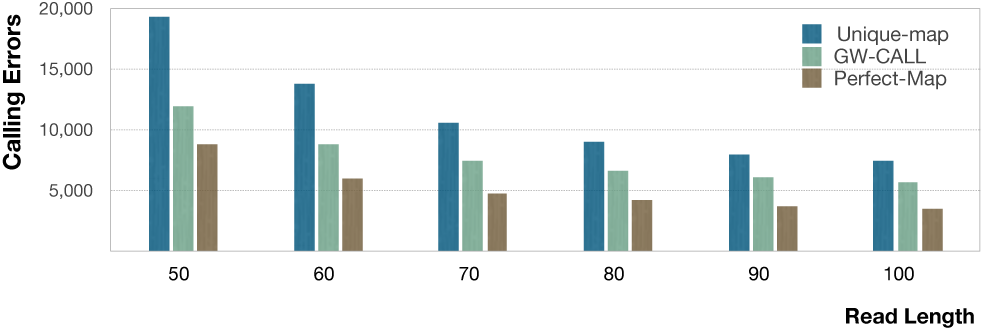
Comparison between different callers for variable read lengths. The other parameters are as follows *p* = 0.01, *c* = 20, and *ϵ* = 0.001.

In Figure 4, we have compared the three variant callers for different coverage depth ranging from 5 to 20 with step size 5. Other parameters are fixed with values: *p* = 0.01, *L* = 70, and *ϵ* = 0.001. The result shows that GW-CALL follows the optimum caller for the given range of parameter and outperforms the unique-map caller.

**Figure 4:**
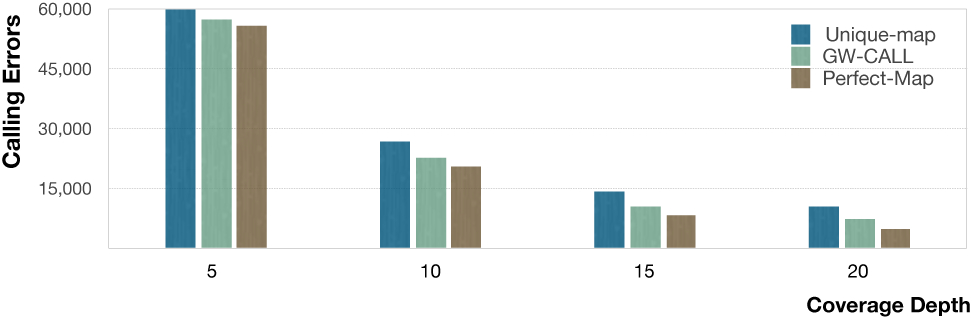
Comparison between different callers for variable coverage depths. The other parameters are as follows *p* = 0.01, *L* = 70, and *ϵ* = 0.001.

In Figure 5, the performance of all three callers is compared for various SNV probability ranging from 0.001 to 0.004. Other parameters are fixed with values *p* = 0.01, *L* = 70, and *c* = 20. The result shows that GW-CALL again outperforms the unique-map caller in all cases, however, its performance degrades substantially as the SNP probability increases. This is an indicator for the opportunity in improving the current version of GW-CALL in case of distance reference and target genomes.

**Figure 5:**
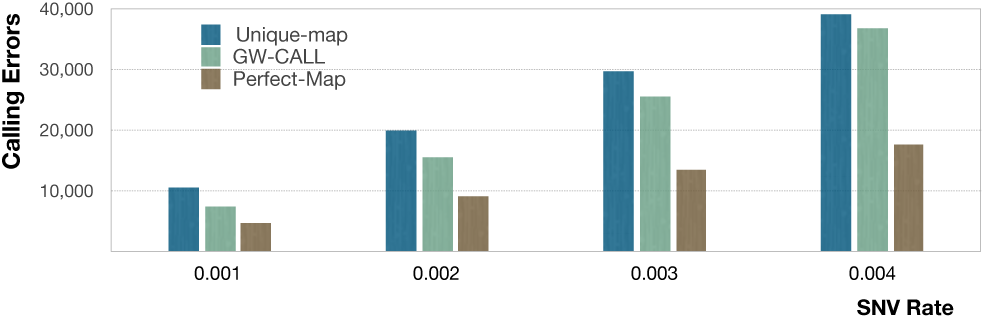
Comparison between different callers for variable SNV rates. The other parameters are as follows *p* = 0.01, *L* = 70, and *c* = 20.

## 4 Discussion

We proposed GW-CALL, a Genome-Wide variant CALLer, that calls variants within less ambiguous genomic loci quickly in the first step and processes the remaining ambiguous locations, called critical points, in a more computationally expensive procedure. Critical points, which cover a very small portion of the genome, mostly lie on the repeat regions where read alignment and variant calling are not straightforward. In fact, local decision making is not enough to capture the complexity of these regions and a global decision making is required. We developed an efficient algorithm that makes a global decision by calling these points collectively.

We showed that our variant caller outperforms the existing methods, especially in redundant genomic regions. Our caller is able to call SNVs given a reference genome and a set of single-end reads with substitution errors. One significant extension is to incorporate mate-pair and paired-end reads in the model. Obviously it can improve the performance of variant calling, since the mapping ambiguity of reads drops greatly by making use of the information of paired reads. Another suggestion is to consider short indel mutations and structural variations in our model. Moreover, we can include indel errors in our read model in order to adapt to situations where sequencing platforms produce reads with indel errors.

## References

[1] Srinivas M Aji and Robert J McEliece. The generalized distributive law. IEEE transactions on Information Theory, 46(2):325–343, 2000.

[2] Kristian Cibulskis, Michael S Lawrence, Scott L Carter, Andrey Sivachenko, David Jaffe, Carrie Sougnez, Stacey Gabriel, Matthew Meyerson, Eric S Lander, and Gad Getz. Sensitive detection of somatic point mutations in impure and heterogeneous cancer samples. Nature biotechnology, 31(3):213–219, 2013.

[3] Kristal Curtis, Ameet Talwalkar, Matei Zaharia, Armando Fox, and David A Patterson. Siren: Leveraging similar regions for efficient & accurate variant calling. Technical report, Tech. rep. UCB/EECS-2015-159. University of California, Berkeley, 2015.

[4] Philippe Galinier, Michel Habib, and Christophe Paul. Chordal graphs and their clique graphs. In International Workshop on Graph-Theoretic Concepts in Computer Science, pp. 358–371. Springer, 1995.

[5] Cyriac Kandoth, Michael D McLellan, Fabio Vandin, Kai Ye, Beifang Niu, Charles Lu, Mingchao Xie, Qunyuan Zhang, Joshua F McMichael, Matthew A Wyczalkowski, et al. Mutational landscape and significance across 12 major cancer types. Nature, 502(7471):333–339, 2013.

[6] Daniel C Koboldt, Ken Chen, Todd Wylie, David E Larson, Michael D McLellan, Elaine R Mardis, George M Weinstock, Richard K Wilson, and Li Ding. Varscan: variant detection in massively parallel sequencing of individual and pooled samples. Bioinformatics, 25(17):2283–2285, 2009.

[7] Daniel C Koboldt, Qunyuan Zhang, David E Larson, Dong Shen, Michael D McLellan, Ling Lin, Christopher A Miller, Elaine R Mardis, Li Ding, and Richard K Wilson. Varscan 2: somatic mutation and copy number alteration discovery in cancer by exome sequencing. Genome research, 22(3):568–576, 2012.

[8] Ben Langmead, Cole Trapnell, Mihai Pop, and Steven L Salzberg. Ultrafast and memory-efficient alignment of short dna sequences to the human genome. Genome Biology, 10(3), 2009.

[9] Michael S Lawrence, Petar Stojanov, Paz Polak, Gregory V Kryukov, Kristian Cibulskis, Andrey Sivachenko, Scott L Carter, Chip Stewart, Craig H Mermel, Steven A Roberts, et al. Mutational heterogeneity in cancer and the search for new cancer-associated genes. Nature, 499(7457):214–218, 2013.

[10] Heng Li. A statistical framework for snp calling, mutation discovery, association mapping and population genetical parameter estimation from sequencing data. Bioinformatics, 27(21):2987–2993, 2011.

[11] Heng Li et al. The sequence alignment/map format and samtools. Bioinformatics, 25(16):2078–2079, 2009.

[12] Heng Li, Jue Ruan, and Richard Durbin. Mapping short dna sequencing reads and calling variants using mapping quality scores. Genome research, 18(11):1851–1858, 2008.

[13] Ruiqiang Li, Yingrui Li, Xiaodong Fang, Huanming Yang, Jian Wang, Karsten Kristiansen, and Jun Wang. Snp detection for massively parallel whole-genome resequencing. Genome research, 19(6):1124–1132, 2009.

[14] Aaron McKenna et al. The genome analysis toolkit: A mapreduce framework for analyzing next-generation dna sequencing data. Genome Research, 20:1297–1303, 2010.

[15] Daniel F Simola and Junhyong Kim. Sniper: improved snp discovery by multiply mapping deep sequenced reads. Genome biology, 12(6):1, 2011.

[16] Robert E Tarjan and Mihalis Yannakakis. Simple linear-time algorithms to test chordality of graphs, test acyclicity of hypergraphs, and selectively reduce acyclic hypergraphs. SIAM Journal on computing, 13(3):566–579, 1984.

[17] Todd J. Treangen and Steven L. Salzberg. Repetitive dna and next-generation sequencing: computational challenges and solutions. Nature Reviews Genetics, 13:36–46, 2012.

[18] Todd J Treangen and Steven L Salzberg. Repetitive dna and next-generation sequencing: computational challenges and solutions. Nature Reviews Genetics, 13(1):36–46, 2012.

[19] Andrew Viterbi. Error bounds for convolutional codes and an asymptotically optimum decoding algorithm. IEEE transactions on Information Theory, 13(2):260–269, 1967.

